# Parallel feedback processing for voluntary control: knowledge of spinal feedback in motor cortex

**DOI:** 10.1101/2024.05.12.593756

**Authors:** Hui Guang, Joseph Y. Nashed, J. Andrew Pruszynski, Hari Teja Kalidindi, Frederic Crevecoeur, Kevin P. Cross, Gunnar Blohm, Stephen H. Scott

## Abstract

How hierarchical feedback pathways coordinate goal-directed motor corrections remains unknown. We show that systematic increases in spinal reflex responses to applied loads lead to a reciprocal reduction in the neural response in primary motor cortex. We use a neural network model to highlight how knowledge on lower-level feedback processing (i.e. spinal reflexes) can be embedded into a network to generate goal-directed corrections through parallel feedback pathways.

## Main Text

Athletic sports like basketball highlight the impressive ability of individuals to shoot with accuracy even if physically bumped by an opponent. A fundamental problem in sensorimotor neuroscience is to understand how such goal-directed motor corrections are generated by highly distributed neural circuits^1-5^. In particular, spinal and cortical feedback both contribute to online corrections within 100ms post-perturbation^6,7^. Each feedback pathway is influenced by various factors and contexts, and it is not clear how these feedback pathways work together to generate goal-directed motor corrections (Fig. 1a).

**Fig. 1.**
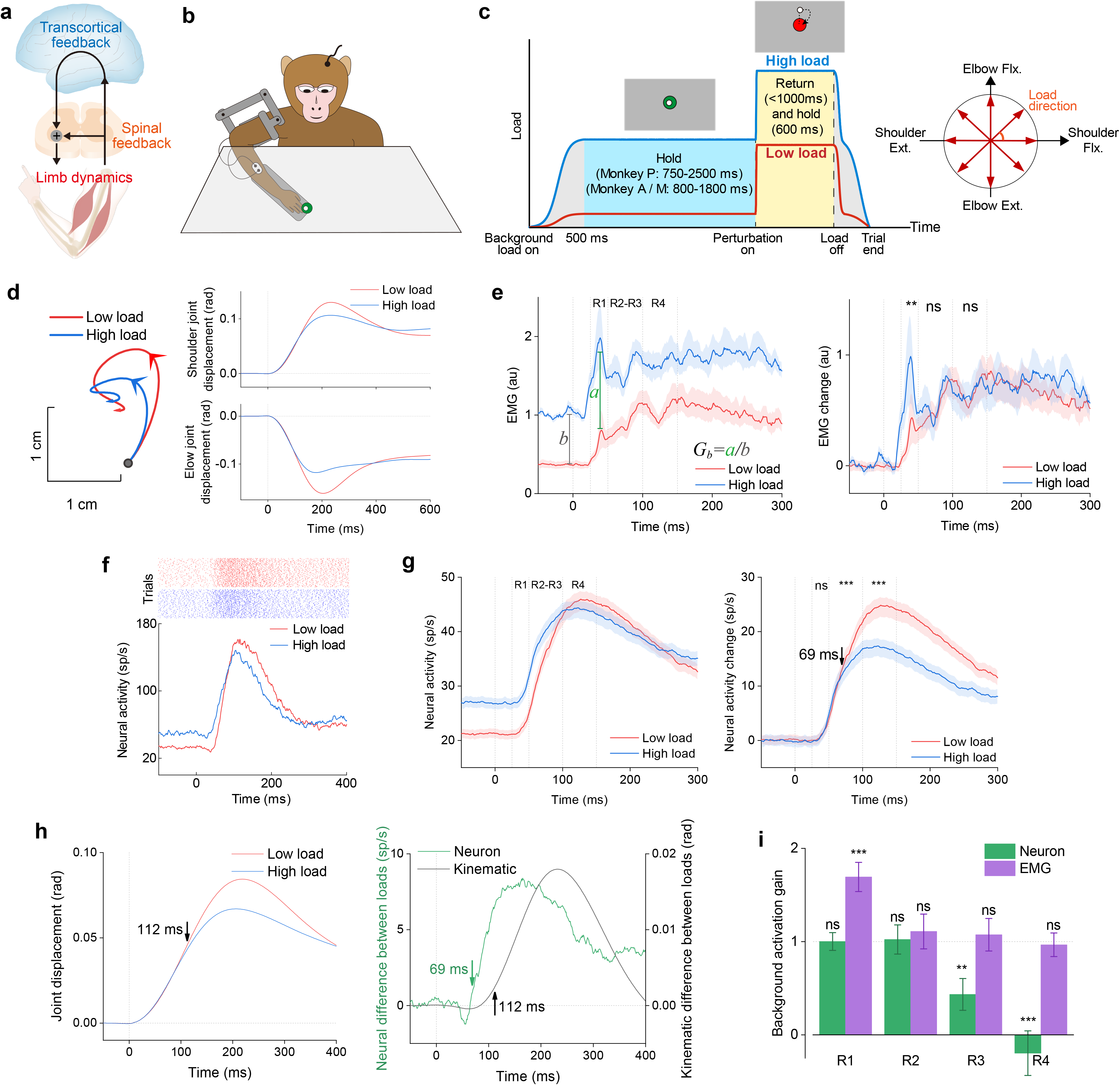
Transcortical feedback response decreases to compensate for increased spinal feedback response. **a**, Hierarchical neuromotor control. Goal-directed motor actions involve parallel feedback pathways including spinal and transcortical. **b**, Experimental setup. Robotic arm applies mechanical loads to the monkey’s arm while performing a posture perturbation task as muscle EMG and neural activity in primary motor cortex are recorded. **c**, The background and perturbation load during the task. The background load with different magnitudes across conditions is initially applied. After a random time period, a step increase in the load was applied. **d**, Hand trajectories and joint displacement in an example load direction for high and low load conditions, where time 0 denotes perturbation onset. Data pooled across monkeys. **e**, Muscle EMGs (±s.e.m) are averaged for the preferred load direction of each muscle and normalized by the baseline activity in the high load condition. R1: 25-50 ms; R2: 50-75 ms; R3: 75-100 ms; R4: 100-150 ms. ** *P*<0.01, paired-sample *t*-test for the significant difference between two loads. **f**, An example neuron from monkey A during the perturbation correction. **g**, Population activity (±s.e.m) for all neurons for the preferred load direction, pooled across monkeys. *** *P*<0.001, paired-sample *t*-test for the significant difference between two loads. Arrow denotes time when neural activity across background load conditions. **h**, Averaged shoulder and elbow joint displacement across load directions. Arrow denotes time when kinematics differ by 5% between the two conditions. **i**, The background load activation gain *G*_*bb*_ (±s.e.m) across epochs. The gain is calculated by the difference in activity between the two load conditions for a given epoch divided by the difference in activity for the background load epoch (Fig. 1e). ** *P*<0.01, *** *P*<0.001, *t*-test for significant difference compared to 1.

The fundamental question we address here is whether variations in spinal reflexes impact cortical feedback responses. We exploited the fact that the spinal stretch reflex increases with background muscle activity, termed gain scaling^8-10^. Three monkeys were trained to perform a posture perturbation task (Fig. 1b). Each trial was initiated by applying a high or low background load and displaying a spatial target. After the monkey maintained its hand at the target for a random time, a step increase in load was applied to the arm, and monkeys had to return their hand to the target within 1000ms to receive a water reward (Fig. 1c). Trials with two background loads combined with eight load directions were presented in random order.

Background load significantly modulated behavior and EMG responses. Across all eight load directions, the monkeys consistently displayed faster corrective responses with smaller displacements following the disturbance for the high versus the low background load (p<0.001, two-way ANOVA Fig. 1d, Extended Data Fig. 1). As previously observed in humans^10^, the spinal stretch reflex from 25-50ms post-perturbation was enhanced for the high versus low background, indicated by a significant increase in perturbation-related EMG during the short-latency epoch (p=0.002, paired sample t-test, Fig. 1e).

Does M1 account for changes in the spinal stretch reflex for high vs. low background load conditions? Critically, we found that while short-latency muscle activity was increased, the perturbation-related M1 activity was reduced for the higher background load. We recorded the activity of 130, 58, and 92 neurons in monkey M, A, and P respectively in the postural perturbation task; only neurons that significantly responded to the perturbation were included (p<0.05, two-way ANOVA). Fig. 1f,g illustrate a common pattern of activity where the background discharge was increased by the background load, but the perturbation response was reduced in size. We averaged neural responses based on the direction with the largest load-related activity (Preferred Direction, PD). We found a significant decrease in the population response between the high versus low background load condition, during the long-latency 50-100ms and 100-150ms epochs (Fig. 1g; p<0.001, paired-sample t-test). Similar reductions were observed for each animal (Extended Data Fig. 2). Notably, a temporal analysis identified a difference at 69ms in M1 responses between background load conditions, whereas kinematic differences were later at 112ms (Fig. 1h). Thus, the reduction of M1 responses was not generated by sensory feedback of limb kinematics. We also found inhibitory responses for loads opposite to a neuron’s PD were smaller for the high background condition (Extended Data Fig. 3).

**Fig. 2.**
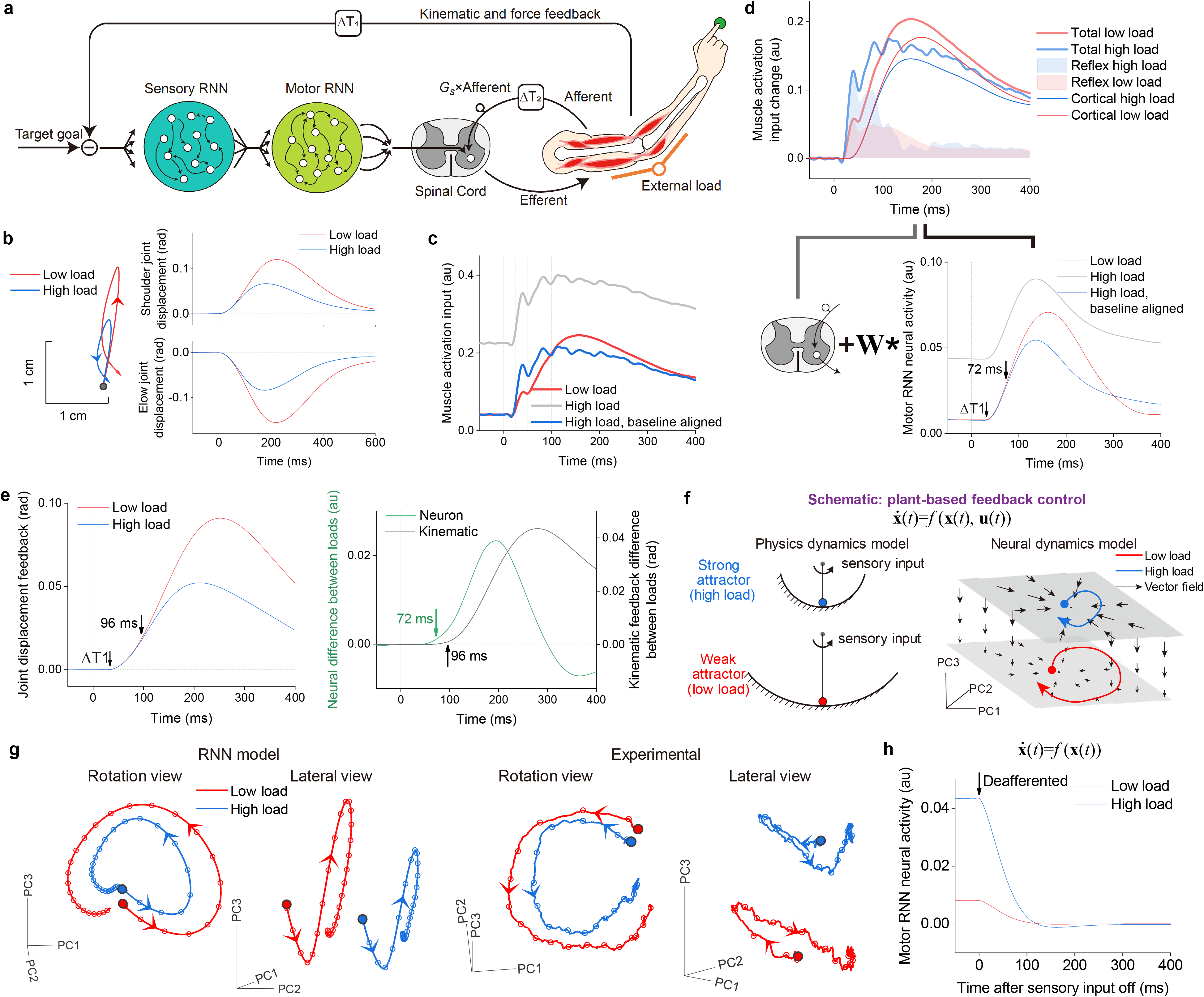
Neural network model exhibits changes in activity due to spinal gain scaling. **a**, The model is composed of recurrent neural networks (RNNs), a spinal reflex pathway, a two-segment musculoskeletal plant and proprioceptive feedback. We set a higher spinal stretch reflex gain *G*_*ss*_ for the high background load condition. **b**, Simulated hand trajectories and joint displacement in the same example direction as Fig. 1d, where time 0 denotes perturbation onset. **c**, Mean muscle activation to agonist muscles. **d**, Output layer RNN population activity and decomposition of muscle activation input into cortical and spinal components. The cortical contribution is computed by projecting output layer RNN neural activities onto muscles via the synaptic weight matrix **W. e**, Averaged shoulder and elbow joint displacement feedback across eight load directions. The moment when neurons differ by 5% between the two conditions was also earlier than observed for the kinematics. **f**, Plant-based feedback control. The RNN can be interpreted as a physical dynamic system, where the neural response is a result of intrinsic dynamics and external sensory input: driven by input but also guided by a vector field at the current state. The vector field is partitioned by initial neural states and varies across neural space, to adapt to changes in the controlled plant and spinal reflexes to generate plant-based feedback control. **g**, Experimental and simulated neural trajectories using principal component analysis in the same load direction Fig. 2b, pooled across monkeys. Each node (open circle) indicates every 20 ms starting from perturbation onset. **h**, Deafferented RNN reveals variations in intrinsic dynamics across neural space. During network testing, the trained RNN is deafferented by setting transcortical feedback gains to 0 at perturbation onset. As a result, the neural response becomes independent of sensory input drive and is solely guided by the vector field derived from initial states. The neural activity exhibits a faster decrease and suggests RNN with a stronger attractor in high load condition.

**Fig. 3.**
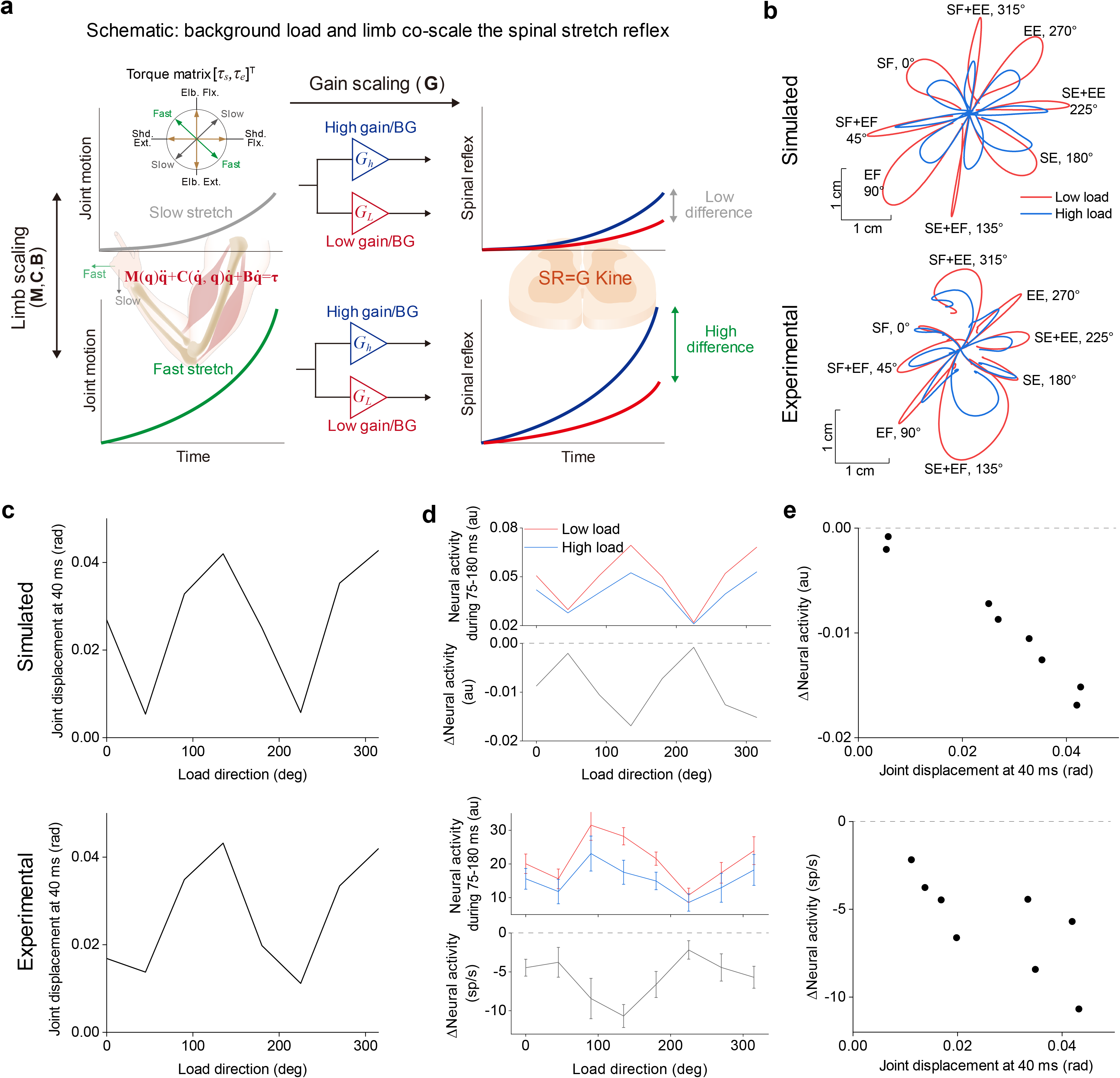
Neural activity in motor cortex accounts for the impact of limb biomechanics on gain scaling. **a**, Schematic diagram highlighting how spinal stretch reflex is influenced by both applied perturbation load and background load. Limb dynamics (inertia, Coriolis, and damping matrices) influence combined flexion load at one joint and extension load at the other (green line) lead to greater joint angular motion, and in turn, an increase in spinal stretch response (left panel). An increase in background load generates a further increase in the spinal stretch response due to gain scaling. **b**, Simulated and actual distribution of hand trajectories across eight load directions. Load directions are defined in joint space as Figs. 1c and 3a. S, shoulder; E, elbow; F, flexion; E, extension. **c**, Tuning curve of joint displacement at 40ms post-perturbation. Load perturbation direction defined in joint-torque space (Figs. 1c and 3a). **d**, Variation in neural activities across load perturbations for high and low background load conditions. Neural activity is calculated from the mean (±s.e.m) during 75-180 ms post-perturbation, and the reduction from low to high load. **e**, Correlations between early joint motion (40ms) and the neural activity difference between high and low load conditions for output layer RNN activity (simulated) and M1 neural activity (experimental).

To quantify if and when background load affected muscle and neuronal activity, we compared the background activation gain (see Methods) of the EMG and M1, illustrated in Fig. 1e where a value above 1 denoted gain scaling^10^. The results demonstrated that the EMG gain during the R1 epoch (25-50ms) was greater than 1 (p=0.001, t-test), and the gains during other epochs were not (p>0.05, Fig. 1i). In contrast, the M1 gain was less than 1 in R3 and R4 epochs (R3, 75-100ms, p=0.002; R4, 100-150ms, p<0.001). Similar patterns were observed for each individual monkey (Extended Data Fig. 4). We also found similar results in a variant of the posture perturbation task with three levels of background load (Extended Data Fig. 5).

To interpret the interactions between neural feedback processes, we created a model composed of recurrent neural networks (RNNs), spinal stretch reflexes, and a two-segment musculoskeletal model of the arm with six muscles spanning the joints, and proprioceptive feedback (Fig. 2a, See Method and ref.^11^). The model was trained to perform the same posture perturbation task, with an optimization function over kinematics and energy costs. We set higher spinal reflex gains for higher background load to test if the RNN can infer this change in spinal reflexes via motor learning.

Consistent with the experimental results, the simulated corrective response was faster with higher background load (Fig. 2b). The muscle activation generated by the combined contributions of spinal and cortical feedback pathways resembled the experimentally observed EMG patterns, with a larger initial R1 response for higher background load, but the difference diminished and became smaller after 100ms (Fig. 2c).

The model enabled the decomposition of system dynamics and separation of spinal and cortical contributions (Fig. 2d). The initial R1 response was generated entirely by spinal feedback with an increased response for higher background load. Notably, activity in the motor RNN responded less to the disturbance for the high background load, and the reduction in the simulated M1 activity was observed at 72ms, which was earlier than kinematic differences observed at 96ms (Fig. 2e). Thus, the RNN controller had embedded knowledge on spinal reflexes cued by the background load. How this embedded knowledge impacts the inherent dynamics in the RNN can be visualized using a vector field representation (Fig. 2f-h). Differences in the initial background load led to a shift in the initial neural state. Consequently, the same kinematic feedback generated when the load was applied leads to different relative patterns of activity in the vector field. This finding was further validated by model variants, demonstrating the feedback control by RNN involved adapting to the influence of spinal feedback and initial state (Extended Data Fig. 6).

Different combinations of joint loads lead to different amounts of joint motion due to limb mechanics^12,13,14^, and thus, influences the size of the spinal stretch response (Fig. 3a-c, Extended Data Fig. 7). For example, flexor torque at one joint and extensor torque at the other lead to larger angular motions than combined flexor torques or extensor torques at both joints. Correspondingly, these differences lead to systematic variations in the M1 response across load directions and background loads, observed in both simulated and experimental data (Fig. 3d, Extended Data Fig. 8). As higher stretch velocity increased the spinal reflex for certain joint load combinations, M1 neurons displayed a further reduction in load-related response from low to high background loads (Fig. 3e, *r*=-0.99, *P*<0.001, simulated; *r*=-0.73, *P*<0.05, experimental). Thus, M1 activity reflects not only information about limb mechanics as observed previously^14^, but also the contribution of spinal feedback pathways.

Knowledge of limb mechanics and spinal reflexes are embedded in RNNs through error feedback learning. Given a physical dynamic system and RNNs, their general forms in state space are both 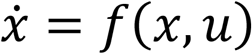. It allows the RNN with sufficient nonlinearity to approximate any physical dynamics, namely the universal approximation theorem^15,16^, and to approximate any nonlinear vector field to control a dynamically complex plant^17,18^. Thus, the RNN was not given direct knowledge of the spinal reflexes, but instead acquired them via inference by feedback error learning cued by background loads^19,20^.

Hierarchical control has previously been proposed in gaze and arm movement models^21,22^. However, how the brain deals with control hierarchies practically might depend on the predictability of the downstream controllers. E.g. spinal cord reflexes are fairly stereotyped and could thus be learned by an upstream controller, as seems to be the case in our study. Conversely, in gaze control, head movement can act as perturbations to the controller, but since head movements are very variable (and under volitional control), it has been suggested that an internal (possibly cerebellar) feedback loop is needed to account for different voluntary head contributions to gaze shifts in addition to accounting for the vestibulocular reflex^21^. Thus, different neural implementations of hierarchical control might exist depending on the system properties.

The present results provide a different perspective from the classic muscles versus movement debate of M1 function^23^. The idea that M1 represents/controls muscles highlights the substantive size of the corticospinal tract including corticomotoneurons that synapse directly onto motoneurons. However, this perspective ignores subcortical and spinal contributions to motor output. Alternatively, the idea that M1 controls movements is captured by most hierarchical models that emphasize serial processing^3,22,24^. Motor cortex represents higher level motor features such as the direction of movement that is converted in the spinal cord into the detailed patterns of muscle activity to generate movement. Our model recognizes multiple levels to the motor system, and emphasizes a parallel organization where both cortical and spinal feedback pathways directly contribute to the generation of motor output. From this perspective, the goal of transcortical feedback is to generate goal-directed motor actions by providing a signal that is the difference between the muscle activity to attain a behavioral goal and the contributions already provided by brainstem and spinal pathways.

## Method

### Animals and apparatus

All animal protocols were approved by the Animal Care Committee at Queen’s University. Three male rhesus macaques were trained to sit in a chair with Kinarm Exoskeleton robotic systems attached to their arm permitting motion of the arm in the horizontal workspace^25^ (Kinarm, Kingston, Ontario, Canada). A virtual reality system displayed a spatial goal (radius 0.95 cm) and visual feedback of hand position (white circle, radius 0.25 cm) aligned with the horizontal workspace (Fig. 1b).

The animals performed a postural perturbation task similar to our previous studies^26,27^, except that at the beginning of a trial, the robot applied one of two background loads to the shoulder and/or elbow joints (randomly selected order; ramp-up time 500 ms; 0.1 and 0.8 Nm for Monkey A, 0.1 and 0.65 Nm for Monkey M, 0 and 0.2 Nm for Monkey P). A central target was illuminated (red out of target, green into target) and the monkey was trained to move to and maintain their hand at this spatial target (approximate arm geometry: 35 degrees shoulder flexion and 90 degrees elbow flexion). Once the monkey maintained the hand in the target for a random period (800-1800ms for Monkeys A and M, 750-2500ms for Monkey P), a step load was applied (0.25Nm for Monkeys A and M, 0.24Nm for Monkey P) and the monkey was rewarded by returning to the target within 1000ms and holding for 600ms (Fig. 1c). The step perturbation was always the same sign as the background load. Data was collected in pairs of opposing flexion/extension loads: (1) shoulder flexion only and shoulder extension only, (2) shoulder and elbow flexion, and shoulder and elbow extension, (3) elbow flexion only and elbow extension only, and (4) shoulder flexion with elbow extension, and shoulder extension with elbow flexion. The load pairs were switched each day. Monkey M and A additionally participated in a second experiment in which there were three different background loads (0.1, 0.45, and 0.8Nm for Monkey A, 0.1, 0.375, and 0.65Nm for Monkey M) and the rest of the task remained the same.

To identify the preferred load direction of neurons, monkeys performed the no-background-load perturbation tasks in eight directions as the first block. It enabled individual neurons to be analyzed in the load direction inducing maximum activation.

### Neural, muscle and kinematic recordings

Neurons from Monkeys A and M were recorded by implanting 96-channel Utah Arrays (Blackrock Microsystems, Salt Lake City, USA) into the arm region of primary motor cortex (M1). Monkey P was implanted with a chamber providing access to the shoulder/elbow region of M1, and neurons were recorded using single tungsten microelectrodes (FHC, Bowdoin, USA). Spike waveforms were sampled at 30 kHz and collected using Grapevine Neural Interface Processor (Ripple Neuro, Salt Lake City, USA) for Monkey A and M or Neural Signal Processor (Blackrock Microsystems) for Monkey P, respectively.

Electromyographic (EMG) activity was recorded by intramuscular electrodes. Monkey P was recorded using two electrode wires inserted percutaneously into the belly of each muscle. Monkey M was implanted with a 32-channel chronic EMG system (Link-32, Ripple Neuro). The eight strands were distributed to individual muscles, and four electrodes of each strand were inserted into the muscle belly. EMG signals were acquired by Neural Signal Processor digitized at 1 kHz for Monkey P and Grapevine Neural Interface Processor digitized at 2 kHz for Monkey M, respectively. We recorded brachioradialis, pectoralis major, brachialis, anterior deltoid, middle deltoid from Monkey P, and brachioradialis, brachialis, triceps long and lateral, pectoralis major, anterior deltoid, posterior deltoid, and biceps long from Monkey M.

The shoulder and elbow angles and angular velocity measured by the exoskeleton robot were also collected by the neural processors and digitized at 1 kHz.

Datasets for Monkeys A and M were collected twice with repeat sampling spaced 6 months apart. Each dataset consisted of 4 sessions of data one each for the 4 pairs of flexion/extension tasks. For Monkey P that was recorded by the single electrode, 59 and 8 sets of neural and EMG data were collected respectively.

### Data processing

Spikes were sorted offline manually (Plexon Offline Sorter, Brussels, Belgium), and isolated units were included as neurons for analysis. The neural firing rate was computed by convolving the spike timestamps with a kernel^28^. We implemented a two-way ANOVA with time epoch (two levels: baseline and 0-300 ms post perturbation) and load direction (eight levels) as factors to identify neurons that were sensitive during the no-background load task. Neurons with a significant effect of epoch or load direction were included (*P*<0.05). For load-sensitive neurons, we identified their preferred load direction (PD) as the load direction that had the largest activation in the post-perturbation epoch. The individual neuron responses in the PD and reverse PD were denoted as agonist and antagonist neurons, respectively.

EMG signals of Monkey M were bandpass filtered by the Grapevine processor at 15-375 Hz during the recordings. For each EMG strand, we computed the signal difference between each pair of proximal and distal contacts, and the signal difference was then rectified and averaged to represent the muscle activation. For Monkey P, EMG signals were bandpass filtered by a 6^th^ order Butterworth filter at 20∼250 Hz, rectified, and low-pass filtered with 6^th^ order zero-phase lag Butterworth filter at 100 Hz. The load direction producing the maximum EMG baseline in the high background load condition was denoted as the PD of individual muscles. EMGs were normalized by the baseline activity in the high background load condition.

To represent the influence of background load on EMG and neural perturbation responses, we calculated the background activation gain *G*_*b*_, which is given by

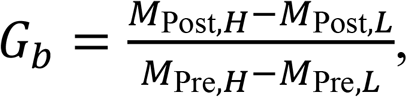

where *M*_Post,*H*_ and *M*_Post,*L*_ denote the mean neural activity or EMG during post perturbation epochs, in high and low background load conditions respectively, and *M*_Pre, *H*_ and *M*_Pre,*L*_ denote the means during pre-perturbation.

### Neural network and limb model

We implemented a neural-limb dynamic model to simulate the neural and kinematic responses during the posture perturbation task. The neural-limb model consisted of three components: recurrent neural networks (RNNs), spinal reflex, and two-segment musculoskeletal model of the arm. Briefly, the RNNs as cortical neural networks received kinematic feedback from the arm and sent descending commands converging on muscles (Fig. 2a). Simultaneously, proprioceptive signals projected to the spinal cord and generated a spinal reflex. The muscles activity reflected the contribution from both cortical input and spinal reflex, and moved a planar limb. The neural-limb dynamics was solved by Euler’s integration with a discretized time step Δ*t* of 10 ms.

The RNNs were composed of two layers to imitate the sensory and motor cortices. The first layer represented the sensory cortex that accepted inputs from sensory feedback, and the second layer imitated the motor cortex as the output layer to generate descending commands to muscles. Each layer consisted of 200 neural units, and the network outputs **y**(*t*) of each layer followed the discretized neural activation dynamics as

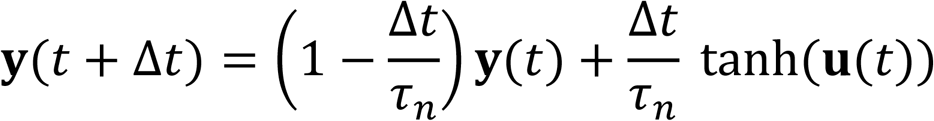

where *τ*_*n*_ of 20 ms denotes the time constant of neural activation dynamics, and **u** (*t*) means the input to each layer.

The inputs to the sensory layer **u**_*s*_ and M1 layer **u**_*m*_ were respectively calculated as

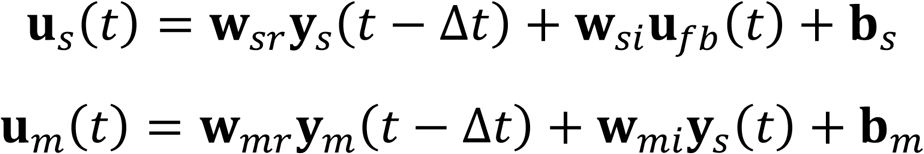

where **y**_*s*_ and **y**_*m*_ are the outputs of sensory and M1 layers respectively, **b**_*s*_, **b**_*m*_, **w**_*si*_, **w**_*mi*_, **w**_*sr*_, **w**_*mr*_ are the bias and weight trained by machine learning, and **u**_*fb*_ denotes the feedback input.

The feedback input **u**_*fb*_ was calculated by the deviation of kinematic states from goal in joint space and muscle force

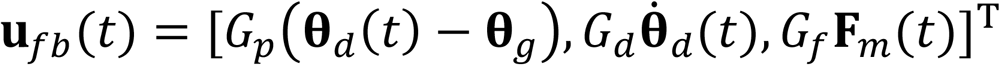

where **θ**_*d*_ denotes the delayed feedback of shoulder and elbow joint angles **θ**_*d*_(*t*) = [*θ*_*s*_(*t* − Δ*T*_1_), *θ*_e_(*t* − Δ*T*_1_)], **θ**_*g*_ indicates the goal joint states at target, and **F**_*m*_ includes the force of six muscles **F**_*m*_ = [*F*_*m*,1_, *F*_*m*,2_, …, *F*_*m*,6_]. The joint feedback was delayed by Δ*T*_1_=40 ms to account for the transcortical afferent delay. *G*_*p*_, *G*_*d*_, and *G*_*f*_ were the feedback gains in position and velocity, and we set all equal to 1.

The output layer projected to muscles to actuate a limb for horizontal planar motion. The limb was constituted by four monoarticular muscles (shoulder flexor and extensor, elbow flexor and extensor), two biarticular muscles (biceps and triceps), and two skeletal segments (humerus, radius/ulna). As the number of M1 neurons was more than number of muscles, we constructed a third-layer neural network, which was denoted as the muscle activation layer, to converge the M1 outputs to muscles. The number of neural units in the muscle activation layer was identical to the number of muscles. Apart from the descending commands from the output layer, the spinal reflex also contributed to the total input **u**_*act*_ to the muscle activation layer. Therefore, the input to the muscle activation layer was given by

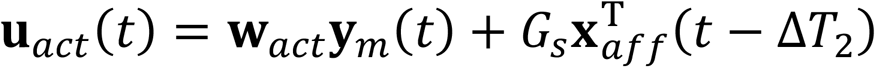

where **w**_*act*_ denotes the synaptic weight, *G*_*s*_ means the stretch reflex gain, and **x**_*aff*_ denotes the afferent. The time delay Δ*T*_2_ of 20 ms accounted for the spinal reflex delay. As the afferent was encoded by muscle spindles that reflected muscle kinematics, the afferent in this study was defined as **x**_*aff*_(*t*) = **G**_*aff*_**x**_*m*_(*t*), where **G**_*aff*_ is a vector of encoding gains, and muscle kinematic state **x**_*m*_ contains the muscle length change from the initial state Δ**x**_*ml*_, stretch velocity **xx**_*mv*_, acceleration **x**_*ma*_, and jerk **x**_*mj*_, namely, **x**_*m*_ = [Δ**x**_*ml*_, **x**_*mv*_, **x**_*ma*_, **x**_*mj*_]^*T*^. Each variable contains the state of six muscles, for example **x**_*mv*_ = [*x*_*mv*,1_, *x*_*mv*,2_, …, *x*_*mv*,6_]^*T*^. As *G*_*s*_ was a scaler, the spinal reflex to each muscle only accounted the afferent from the homogeneous muscle. We set **G**_*aff*_= [1, 0.1, 0.01, 0.0001]. *G*_*s*_ were 0.8 and 1.9 for low and high reflex gains, which was consistent with the experimental data that the EMG value in the high load conditions was ∼2.3 times larger than the low load conditions.

The neural input to the muscle activation layer induced the muscle activation **y**_*act*_, which is given by

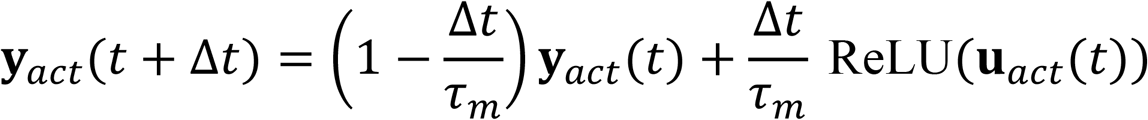

where *τ*_*m*_ of 50ms denotes the time constant of muscle activation dynamics.

The muscle activation was delayed by 30ms and then converted to the contraction force **F**_*m*_ via the muscle force-length (FL) and force-velocity (FV) properties, to produce the joint torques^29,30^. By combining with the external load applied by the robot, the motion of skeletal segments was derived by Lagrange’s equation as a two-segment model. The limb motion recurrently altered the muscle length, which was computed as

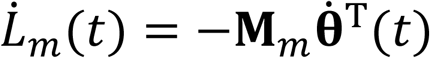

where *L*_*m*_ denotes the muscle length, and **M**_*m*_ is the movement arm of a muscle across the two joints denoted by **θ**(*t*) = [*θ*_*s*_(*t*), *θ*_e_(*t*)]. The changed muscle state updated the reflex via the afferent encoding and muscle force via FL and FV properties.

### Training the neural network for the background load and perturbation task

The objective of training the network is to generate a control policy that can maintain posture against external perturbations corresponding to slowly rising background loads followed by sudden perturbation loads applied in a sequence. We want to derive an ideal actor that acts based on specifying the task goal^11,30^, and not to fit the control policy to the experimentally observed detailed arm trajectories explicitly. This is achieved by formulating a composite cost function that penalized the energy consumption and steady-state kinematic error in the two conditions:

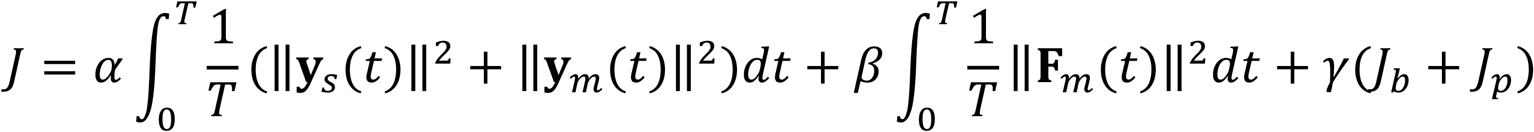

where the first two terms penalize the neural activity (**y**_*s*_ and **y**_*m*_) and force output (**F**_*m*_) throughout the task duration respectively, while the last term is specific to the kinematic cost in background and perturbation loads; *α, β*, and *γ* are the penalty coefficients. Specifically, the kinematic cost is computed as

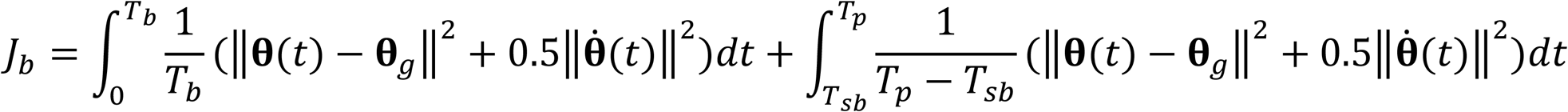

and,

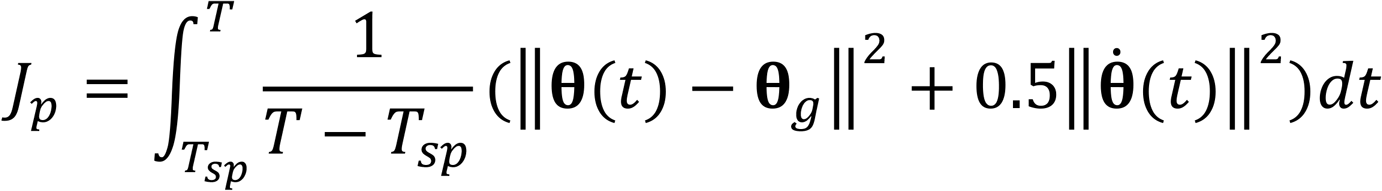

where *T*_*b*_, *T*_*p*_, and *T* denote time of background load onset, perturbation load onset, and total motion; *T*_*sb*_ and *T*_*sp*_ denote the time of steady-state realization of background load and perturbation load. Indeed, as the steady-state time was an artifact value accounting for the kinematic penalization, the actual steady-state time solved by the model was ultimately determined by the model dynamics and the compromised to the energy cost. We set *T*_*b*_ =200ms, *T*_*p*_ =1500ms, *T* =3000ms, *T*_*sb*_ =1000ms, and *T*_*sp*_ =2500ms. Penalty coefficients were defined as *α* =0.0001, *β*=0.01, and *γ*=0.5.

The external torques from the robot with time were the input to the model. Specifically, for the network training, there were 16 load conditions (8 load directions × 2 load sizes) defined in 2 external load matrices (shoulder and elbow), whose columns and rows indicated conditions and time, respectively. The background loads were respectively 0.1 Nm and 0.55 Nm for low and high load conditions, and the perturbation load was 0.25 Nm across all conditions.

The neural network was trained by using the back-propagation algorithm of Adam optimization implemented in the Python and PyTorch machine learning library^31^. Bias **b** was initialized as 0, and weight matrices **w** were initialized by a uniform distribution between 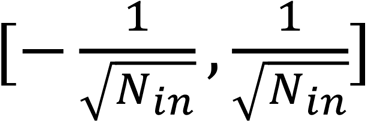, where *N*_*in*_ indicates the number of inputs that each neuron connected in their weight matrix. The training epoch was set as 3000 to ensure a small and converged loss value.

## Acknowledgment

We thank Kim Moore, Simone Appaqaq, Justin Peterson, Helen Bretzke, and Mike Lewis for their laboratory technical assistance; Dr. Douglas Cook for his contribution to monkey surgery and array implant; Dr. Ethan Heming for his support to setup the tasks in Kinarm. This work was supported by the Canadian Institutes of Health Research (CIHR). S.H.S. was supported by a GlaxoSmithKline–Canadian Institutes of Health Research Chair in Neurosciences.

## Author Contributions

Conceptualization, H.G., J.Y.N, J.A.P., F.C., G.B., S.H.S.; Data collection, H.G., J.Y.N, J.A.P., K.P.C.; Data analysis, H.G., J.Y.N.; Simulation, H.G., H.T.K.; Writing manuscript, H.G., S.H.S.; Review and revise manuscript, H.G., J.Y.N, J.A.P., H.T.K., F.C., K.P.C., G.B., S.H.S.; Resources and Supervision, S.H.S.

