## Supplementary Figures for "Parallel feedback processing for voluntary control: knowledge of spinal feedback in motor cortex"

### Extended Data

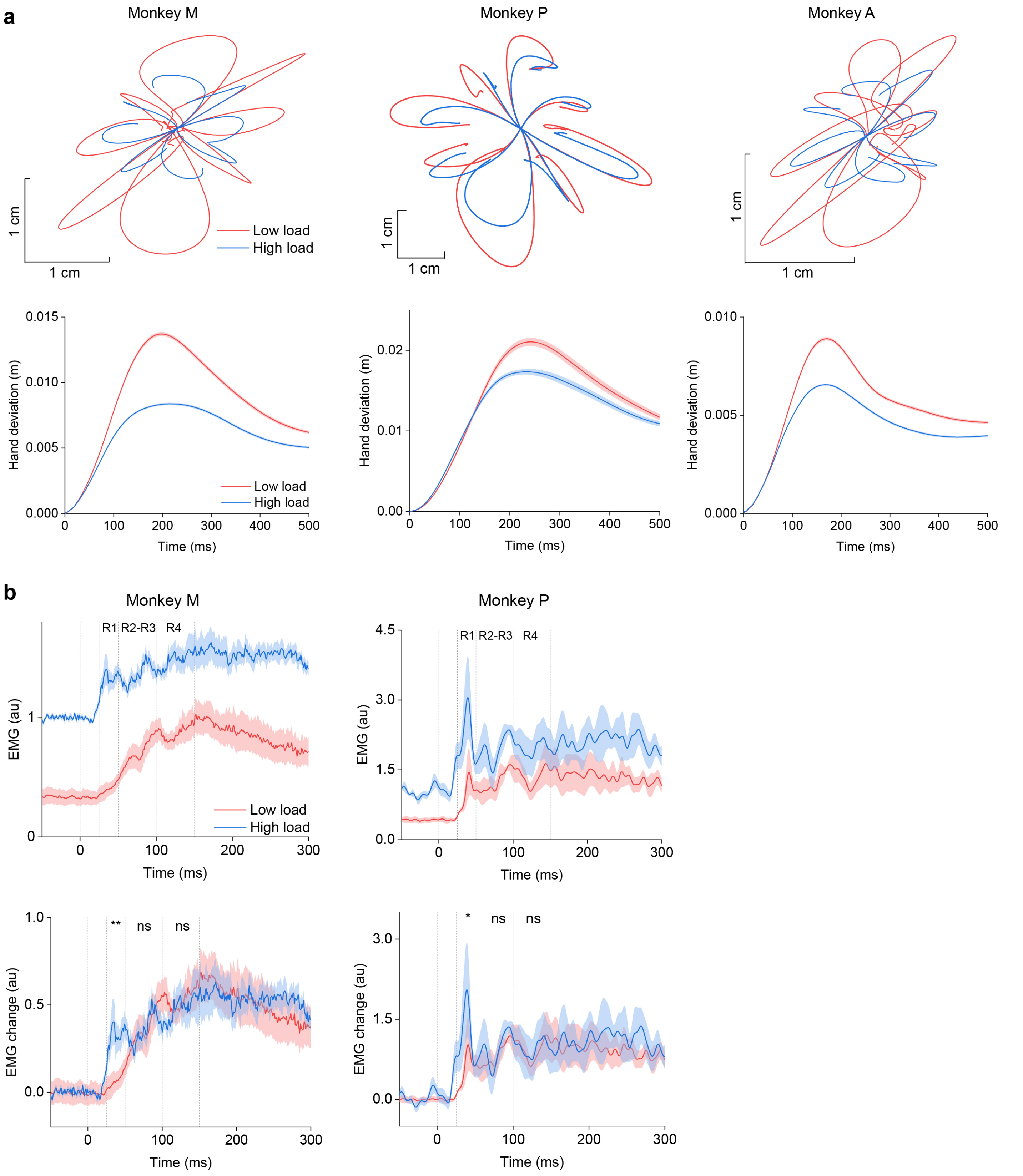


**Extended Data Fig. 1 | Kinematics and EMGs of individual monkeys. a**, Hand trajectories and hand deviation (±s.e.m) from the target post-perturbation. The high background load leads to significant reduction in hand motion compared to the low background load (*P* <0.001, *t*-test, for the peak values). **b**, Population EMG (±s.e.m) of individual monkeys. The EMG of each muscle is for its preferred load direction (load generating maximal activation). The high background load significantly enhances muscle activation to the low background load, during the short latency epoch 25-50 ms representing the spinal reflex (* *P* <0.05, ** *P*<0.01, paired-sample *t*-test).

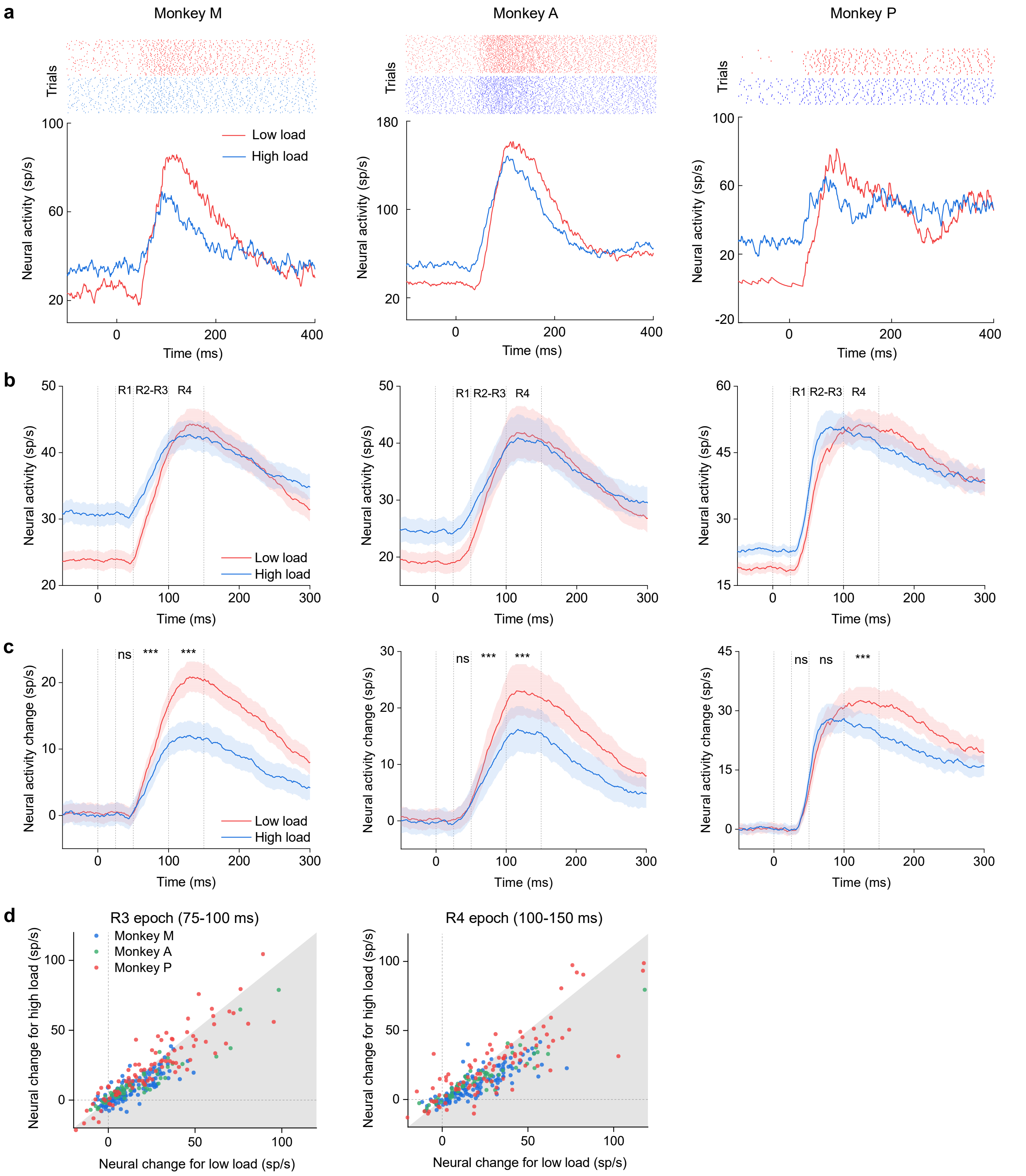


**Extended Data Fig. 2 | An exemplar and population neural activity for individual monkeys. a**, Exemplar neurons. **b**, Population activity in M1. The neural activity is analyzed in the preferred load direction (load that generates maximum activation). **c**, Neural activity response from baseline. The neural response post-perturbation is significantly smaller in the high background load condition compared to the low background load condition, during the R2-R3 epoch (50-100 ms) and R4 epoch (100-150 ms) for Monkey M and A, and during the R4 epoch for Monkey P (*** $P$<0.001, paired-sample $t$-test.). **d**, Individual neuron's change from baseline in high background load condition versus low background load condition, in the R3 (75-100 ms) and R4 (100-150 ms) epochs. The individual neurons within the gray area denote the converse gain scaling effect.

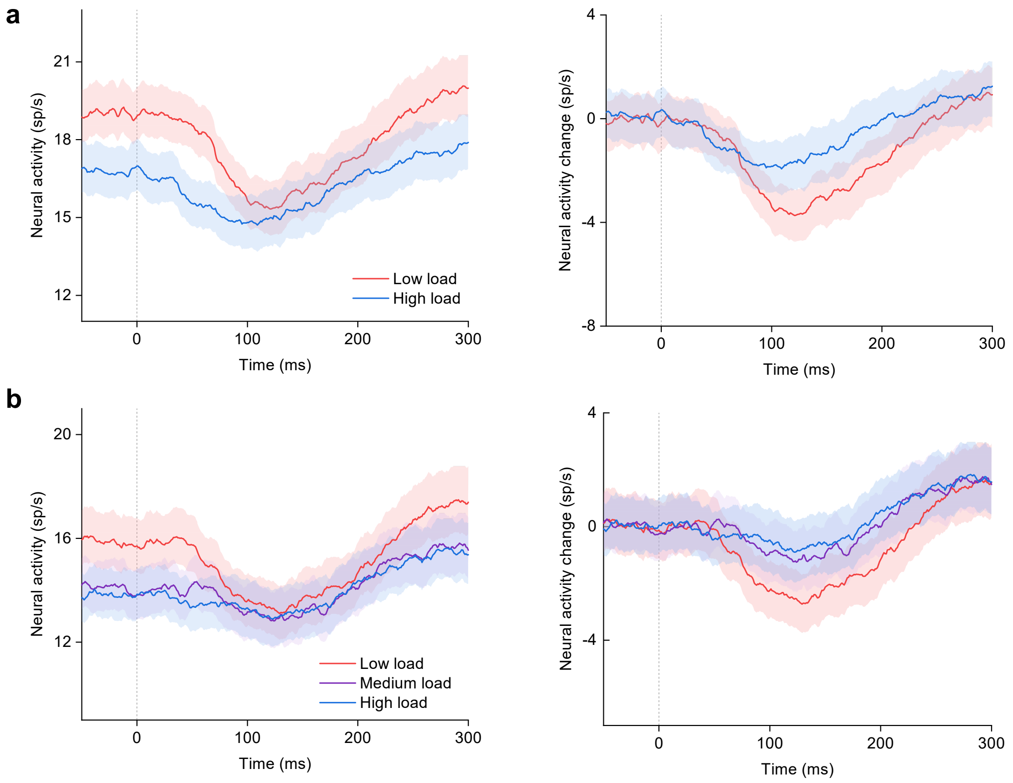


**Extended Data Fig. 3 | Population response post-perturbation in M1 for the load direction opposite to preferred load direction, pooled across Monkey A and M. a**, Two-level background load experiment. **b**, Three-level background load experiment. Left panel denotes population activity (±s.e.m); right panel denotes population activity change from baseline (±s.e.m). The high background load led to a larger reduction in activity post-perturbation as compared to the low background load.

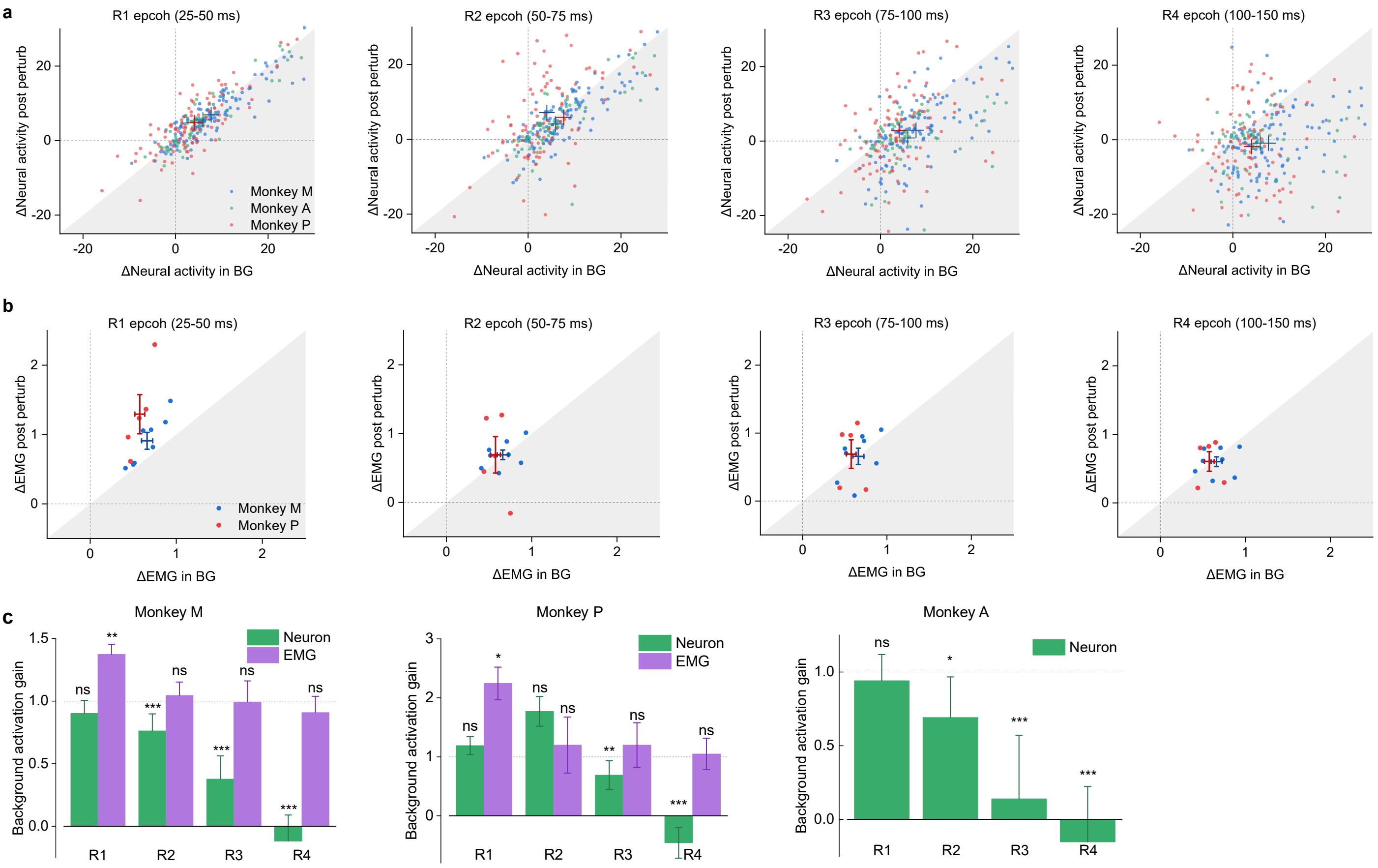


**Extended Data Fig. 4 | Background activation gains of neurons and EMGs. a**, Difference in M1 neural activity between high and low background load conditions during post perturbation epochs vs. background epoch (BG). The data within the gray area denotes the gain less than 1 (gain reduction), and the cross shows the populational mean. R1, 25-50 ms; R2, 50-75 ms; R3, 75-100 ms; R4, 100-150 ms. **b**, Same as **a** but for EMGs. The data within the white area denotes the gain greater than 1 (gain scaling). The cross shows the populational mean with the vertical/horizontal bars denoting the s.e.m. **c**, The background activation gain of populational EMG and M1 neural activity (±s.e.m). The EMG background activation gain is significantly larger than 1 in the R1 epoch (gain scaling), whereas the neural background activation gain is significantly less than 1 for R2 epoch of Monkey M and A, and for R3 epoch of Monkey P, denoting gain reduction (* $P$<0.05, ** $P$<0.01, *** $P$<0.001, $t$-test).

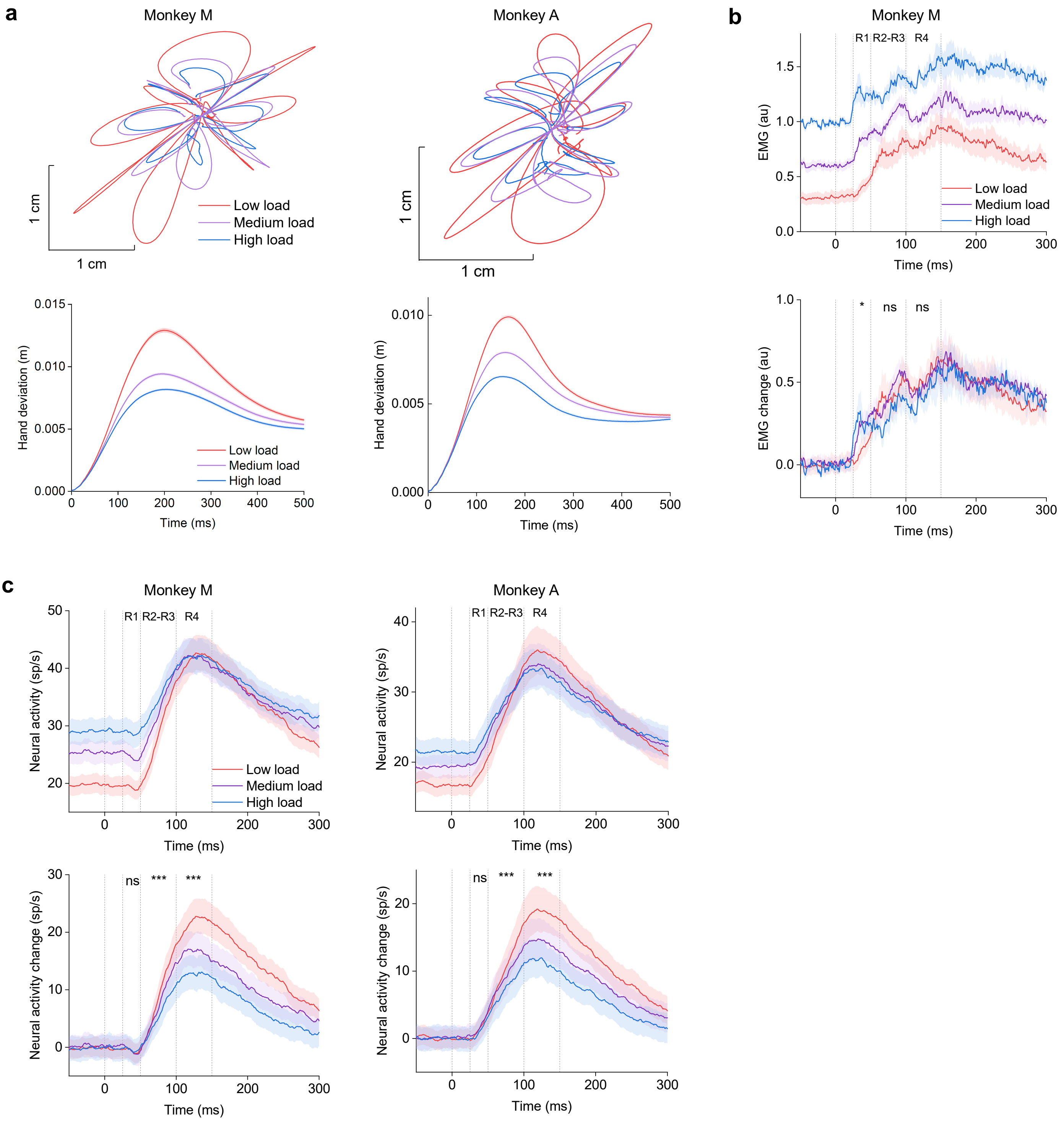


**Extended Data Fig. 5 | Kinematics, EMG, and neurons for three-level background load experiment. a**, Post-perturbation hand trajectories (±s.e.m) for individual monkeys. Higher background load significantly reduces post-perturbation hand motion as compared to the lower background load ($P$<0.001, for peak values between every two load conditions, ANOVA). **b**, Population EMG (±s.e.m) activity of Monkey M. The EMG of each muscle is included for its preferred load direction (load direction generating maximum activation). The high and medium background load led to a significant increase in muscle activation as compared to the low background load, during the short latency epoch 25-50 ms (* $P$<0.05, paired-sample $t$-test). **c**, Population activity for M1 for individual monkeys. Data is analyzed for neuron's preferred load direction (load direction generating maximum activation). The neural activity response from baseline is significantly smaller in the higher background load condition than the lower background load condition ($P$<0.001, between every two load conditions, paired-sample $t$-test), during R2-R3 epoch (50-100 ms) and R4 epoch (100-150 ms).

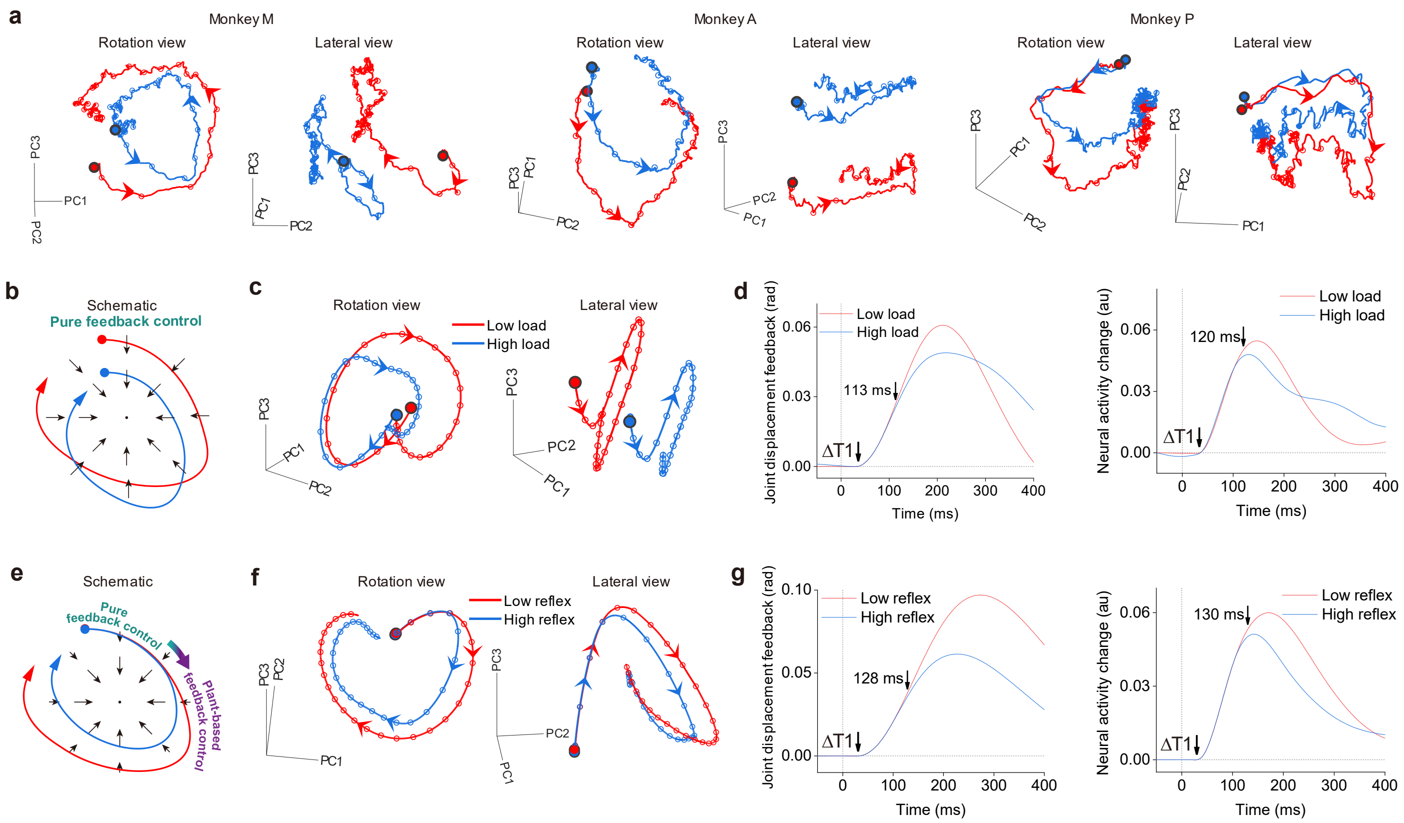


**Extended Data Fig. 6 | The neural trajectories and variants of RNN model. a**, Neural trajectories for individual monkeys using principle component analysis in the same load direction as in Fig. 2g. Each node indicates every 20 ms starting from perturbation onset. **b-d,** A variant of the RNN model in the same load direction. The RNN is trained without incorporating a spinal gain scaling model but is tested with spinal gain scaling. However, the RNN fails to learn the spinal gain scaling, resulting in a delay to 120ms (arrow) in the timing of a 5% neural signal difference between background load conditions, later than kinematic feedback differences (113ms). The vector field, which represents neural dynamics, is not influenced by changes in reflexes within the plant. Consequently, the neural trajectory only differentiates into two curvatures after the sensory input is differentiated (schematic, **b**; simulated, **c**), and it transforms into a pure feedback controller when a new plant with gain scaling modifies the sensory input. We refer to it as a pure feedback controller because the vector field is solely based on the plant learned during training, and it maintains the same mapping between the sensory input and motor command output even when the plant undergoes changes. **e-g,** Another variant of the RNN model in the same load direction as Fig. 2g. Training of high gain and low gain stretch reflex conditions was with a same background load size. In this case, the RNN exhibits a same initial neural state for the two reflex conditions. When driven by the same sensory input, these similar initial neural states evolve into the same neural trajectories, as they are influenced by a common vector field. However, the subsequent gain scaling modifies the post-perturbation motion and sensory feedback, leading the neural trajectories to diverge into different regions of neural space. As a result, the timing of 5% difference between load conditions in neural signals occurred slightly later than difference in joint displacement. The RNN is capable to derive distinct vector fields once the neural trajectories bifurcated, thus enabling control of the plant in the two conditions. These two variants highlight the influence of spinal feedback processing and the need for initial neural states to improve learning in the RNN.

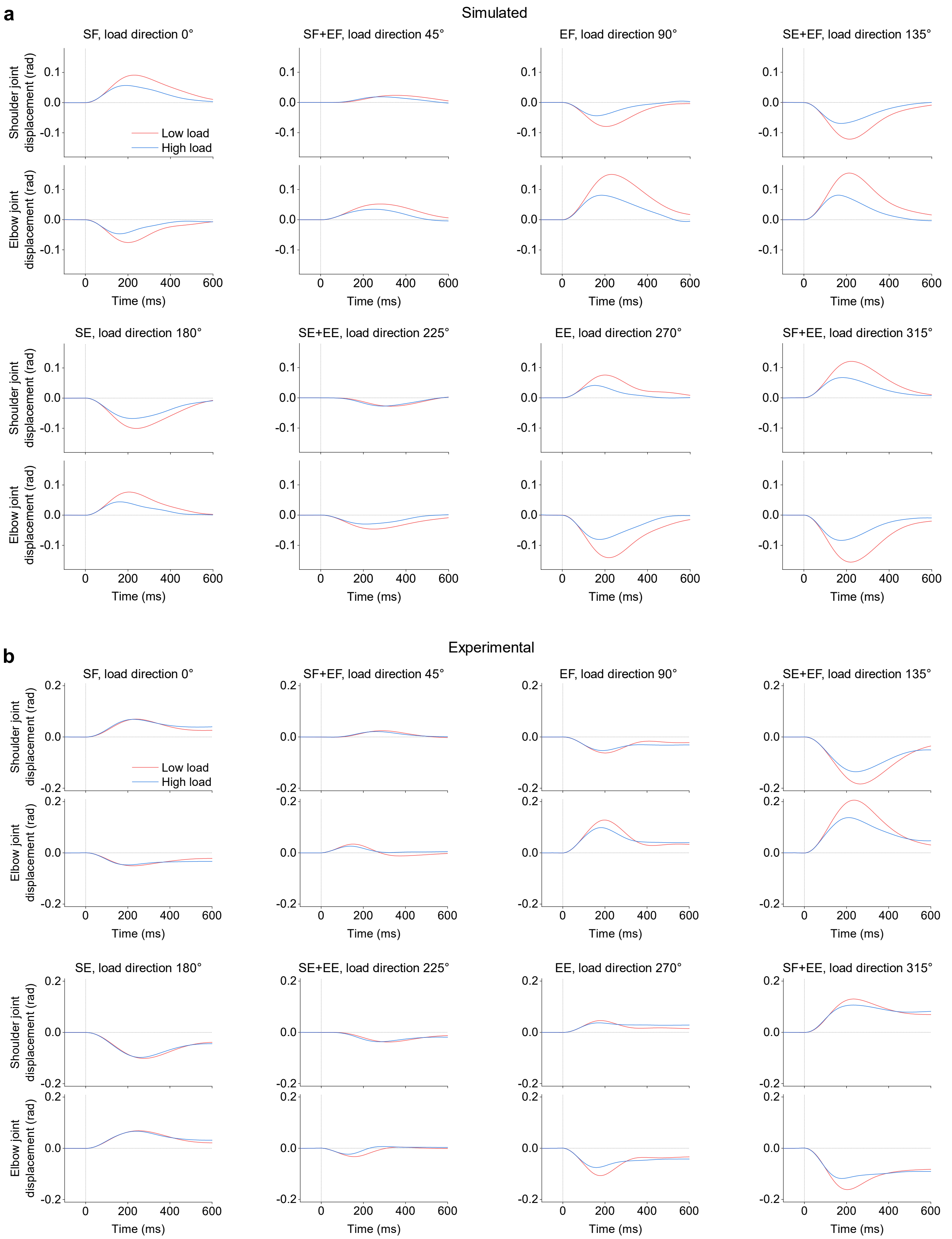


**Extended Data Fig. 7 | Kinematics of joint motion for each load direction. a**, Simulated. **b**, Experimental, pooled across monkeys. Due to limb mechanics, the perturbation uniformly distributed in joint space produces inconsistent joint motion across the eight load directions. In the load direction of SE+EF and SF+EE that drives the limb to move backward and forward, the joint motions are relatively large. Conversely, in the load direction of SF+EF and SE+EE that drives the limb to move leftward and rightward, the joint angular motions are smaller. The larger angular motions enhance the spinal reflex and also the gain scaling effect. S, shoulder; E, elbow; F, flexion; E, extension.

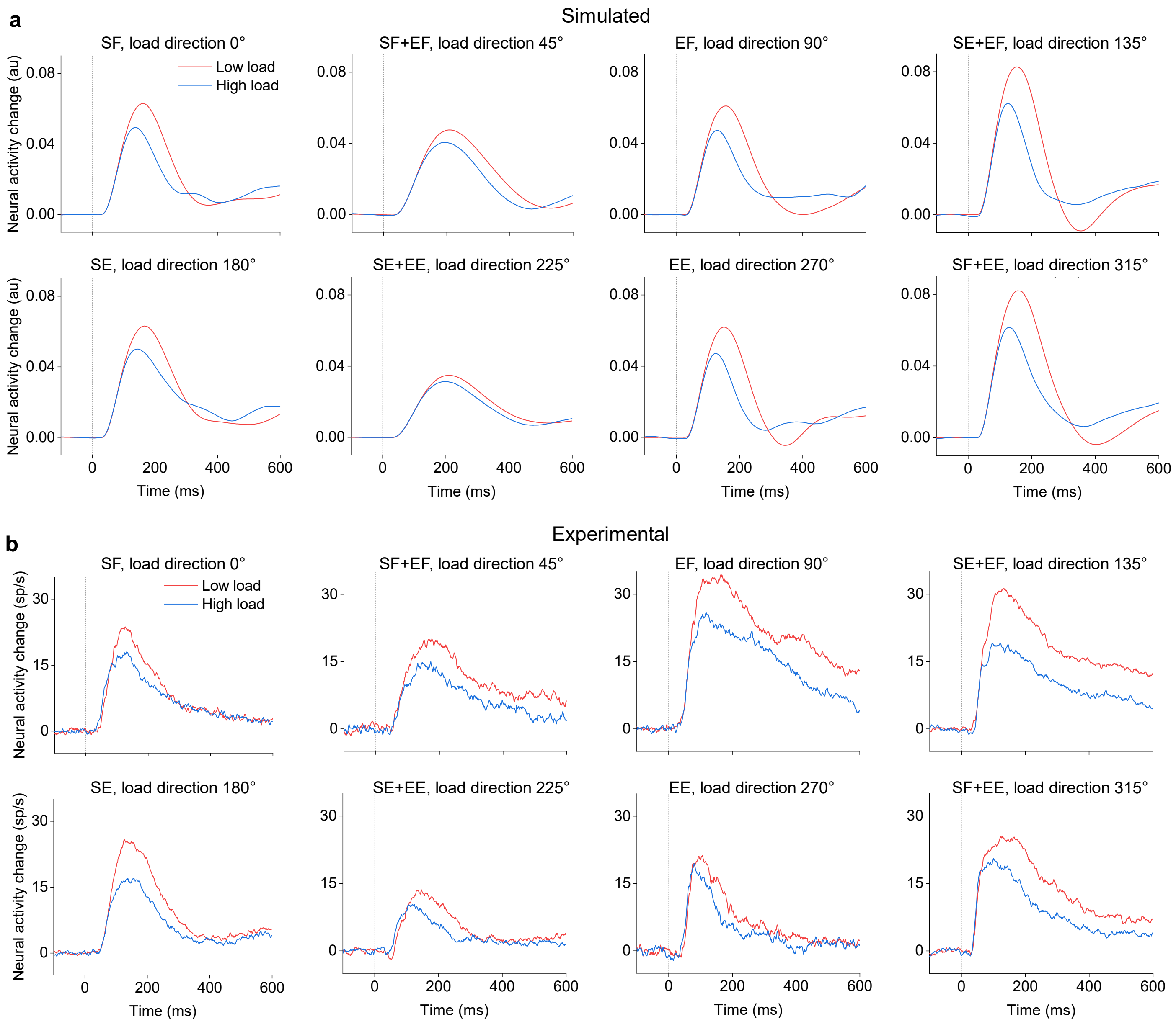


**Extended Data Fig. 8 | Neural responses for each load direction. a**, Simulated. **b**, Experimental, pooled across monkeys. Correspondingly, the neurons generate inconsistent response activities across eight load directions to counter the inconsistent kinematic deviation and gain scaling effect shown in Extended Data Fig. 7. In the load direction with a faster stretch velocity that enhances the gain scaling effect, the neurons show larger responses from baseline and more reduction between high and low background load conditions. It highlights that the M1 activity reflects not only the information about limb mechanics but also how it impacts the contribution of spinal feedback. Namely, the M1 cortex learns the knowledge of spinal reflex integrated with limb mechanics. S, shoulder; E, elbow; F, flexion; E, extension.
